# *Clerodendrum Chinensis Extracts;* AeC and EeC Exerts Rapid Antihypertensive Effects in L-NAME Hypertensive Experimental Models

**DOI:** 10.1101/2020.07.18.210070

**Authors:** Joy I. Odimegwu, Tolulope F. Okanlawon, Obumneme Noel, Ismail Ishola

## Abstract

**Background:** The rise in occurrence of hypertension, a non-communicable disease and a major factor for chronic renal failure, cardiovascular disease, and stroke, which most times lead to sudden death is worrisome. Resistant hypertension is more common and may have no symptoms at all for months or years, but then can cause heart attack, stroke, and vision and kidney damage. Prevention and quick management of hypertension are therefore essential in reducing the risk of these debilitating ailments. Aqueous and ethanolic extracts of the leaves of *Clerodendrum chinensi*s (**AeC** and **EeC**) are used by local communities of West Africa as medicine for rapid antihypertensive actions. We aim to discover the scientific basis for the use of the herb as medicine.

**Methods:** This work investigates the antihypertensive effects of **AeC** and **EeC** in L-Arginine Methyl Ester Hydrochloride (L-NAME)-induced hypertensive rats Acetylcholine, L-Arginine and Sodium Nitroprusside were used as standards. All results were expressed as means ± standard error of mean. Differences were considered significant at p <0.05.

**Results:** Intravenous administration of the extracts caused a significant decrease in the Mean Arterial Blood Pressure (MABP) in a dose-dependent manner. **AeC** at 100mg/kg caused a significant decline in blood pressure in a dose-related manner. Likewise at 100mg/kg, **EeC** reduced MABP steadily from 103.9± 2.55 to 34.1± 0.95mmHg. The two extracts; possess significant antihypertensive properties.

**Conclusions:** Both extracts show significant antihypertensive effects and at high doses could lead to hypotension and so should be used with care. Further research is necessary to determine safe dosage forms.

## 1.0 INTRODUCTION

Cardiovascular diseases (CVDs) actually encompass several coronary artery diseases (CAD) like Angina, Myocardial infarction among others (Mendis *et al.*, 2011). Other CVDs are cardiomyopathy, hypertension, rheumatic heart disease, heart arrhythmia, peripheral artery disease, and venous thrombosis (Mendis *et al.*, 2011). These diseases are being witnessed in larger percentage of the population and in younger subjects too. Also patients have reported a certain resistance of the disease conditions to known drugs.

CVDs are the highest cause of deaths all over the world. WHO, in 2008 reported that more than 17 million people died from CVDs complications that year alone of which more than 3 million of these could have been prevented by changes in diet and lifestyle. Hypertension, the most common CVD also known as high blood pressure, is a medical condition in which the blood pressure in the arteries is persistently elevated (Naish and Court, 2014). There has been an increase in incidences of hypertension with the proportion of the global burden of disease attributable to hypertension increasing significantly from 4.5% in 2000 to 7% in 2010 (WHO, 2008). Hypertension leads to complications with considerable morbidity and mortality which is responsible for at least 45% of deaths due to heart disease and 51% of deaths due to stroke (WHO, 2008). The number of people with uncontrolled hypertension has increased to about 1 billion worldwide in the past 30 years (Danaei *et al.*, 2011). The prevalence of hypertension cases is increasing worldwide; in bothe developed and the developing worlds and seen more and more frequently in younger patients (D’Agostino *et al.*, 2008). Sustained high blood pressure is considered responsible for many serious medical conditions such as arteriosclerosis, heart failure, kidney failure, blindness, and cognitive impairment, and also for many deaths from stroke and heart disease. Systolic, diastolic, and pulse pressure are important predictors of cardiovascular risk (Benetos *et al.*, 2002)

Scientists are reverting to nature for solutions to ailments hence the renewed great interest in plants with antihypertensive compounds used locally but not certified scientifically. *Clerodendrum chinensis* [Osbeck] Mabb. Figure 1 (Chinese glory bower, glory tree, Honolulu rose, stick bush) is a species of flowering plant in the genus *Clerodendrum*. It is placed in the family Lamiaceae, a native to Asian countries of China, Korea, Taiwan, Japan, Vietnam, Western Samoa and American Samoa (PROSEA, 2012) where it grows commonly as a weed along roadsides and as an ornamental shrub with stout branches. The plant has several synonyms; *C. fragrans*, *C. japonicum* etc (Roskov et al., 2019). The leaves’ tinctures are diuretic (Nguyen, 1993). A decoction of the leaves is used in the treatment of blenorrhoea (Chlamydial conjunctivitis) and reputedly a remedy for difficult cases of scabies. Juice from the leaves is an ingredient of herbal bath for children with furuncles (PROSEA, 2012). The plant is used for the treatment of rheumatism and ague, and as an ingredient of a mixture for treating skin problems (De Filipps, 2004). The root extracts are considered antibacterial, and diuretic. It is used in the treatment of abdominal pain, intestinal disease and kidney dysfunctions. It is said to have been used successfully in the treatment of jaundice, lumbago, and hypertension (PROSEA 2012). It is often grown as an ornamental, the double-flowered but sterile form being most commonly cultivated (Valkenburg and Bunyapraphatsara, 2001).

**Figure 1.**
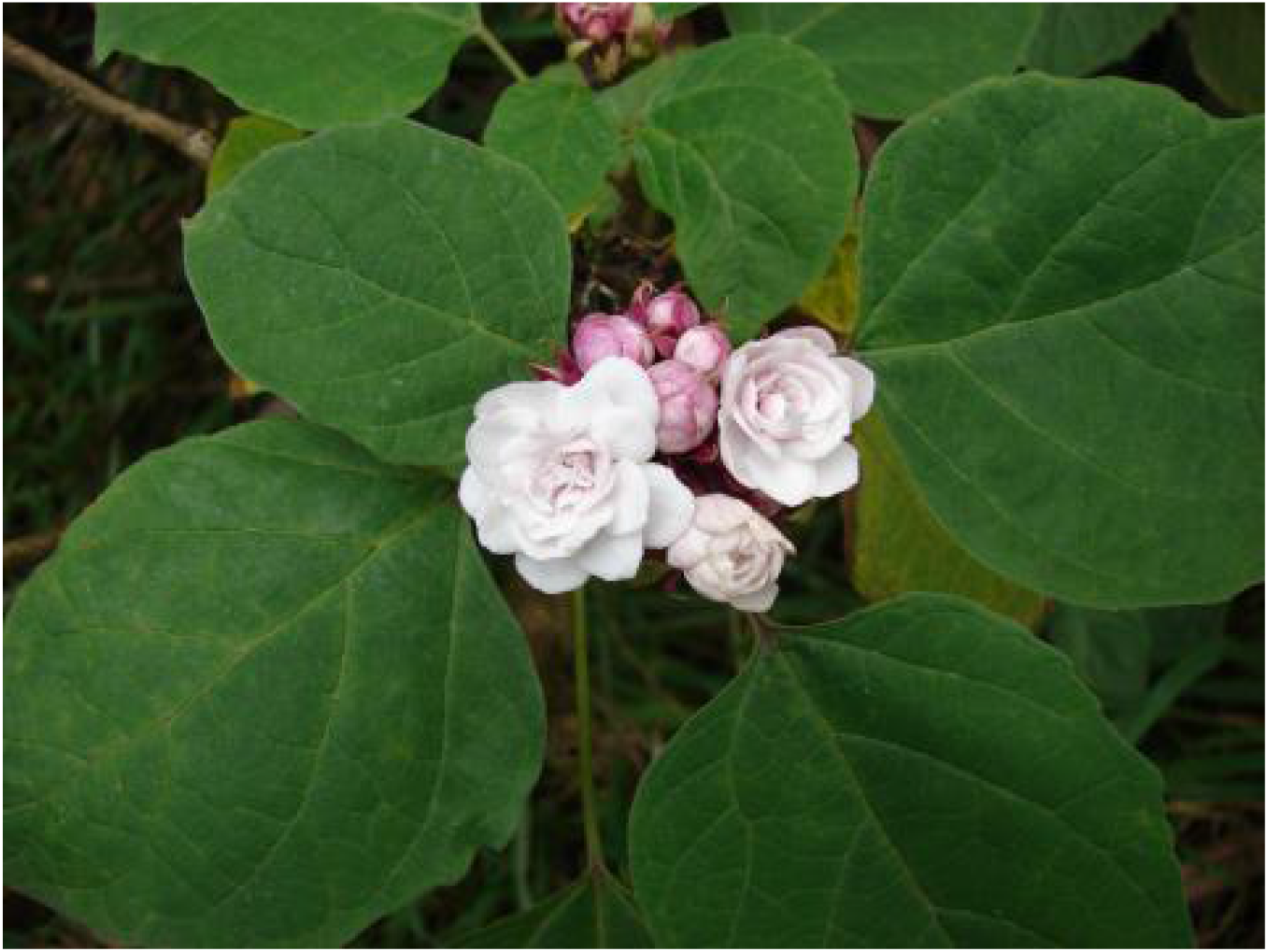
Leaves and flowers of *C. chinensis* [From Tropical Plants Database]

Chemicals frequently employed in hypertension treatment includes; Sodium nitroprusside (SNP), a water-soluble sodium salt comprised of Iron (Fe2+) complexed with nitric oxide (NO) and five cyanide anions. In the living system, it functions as a pro-drug, reacting with sulfhydryl groups on erythrocytes, albumin, and other proteins to release Nitric Oxide (Ribeiro *et al.*, 1992 and Razny *et al.*, 2011). Nitric Oxide, or endothelium-derived relaxing factor, stimulates guanyl cyclase to produce cyclic GMP, sequestering calcium and inhibiting cellular contraction. At the tissue level, these effects of Nitric Oxide result in reduced vascular tone in muscular conduit arteries. Nitric Oxide released from nitroprusside decreases cerebral vascular resistance (Bunbupha, *et al.*, 2015).

L-arginine (L-ARG) is an amino acid with lipid peroxidation-lowering effects which may account for its beneficial effect on Systolic Blood Pressure (SBP) and vascular responsiveness (Ozçelikay *et al.*, 2000). The objectives of this work are the study the antihypertensive effects of **AeC** and **EeC** and evaluate its safety by acute toxicity of using L-NAME induced hypertensive rats.

## 2.0 MATERIALS AND METHODS

L-NAME (Sigma Aldrich, USA). Heparin, Acetylcholine and Sodium Nitroprusside from the Department of Physiology, College of Medicine University of Lagos, Nigeria. L-Arginine was obtained from the Department of Pharmacology, College of Medicine University of Lagos. The leaves of *Clerodendrum chinensis* were originally from plants collected from Abidjan, Ivory Coast and cultivated in a private garden at Lekki, Lagos State, Nigeria 6°29′36″N 3°43′14″E and collected for drying and processing. Verification of the plant identity was done at the Herbarium of the Department of Botany, University of Lagos, Nigeria and assigned voucher number FIH 6974.

### 2.1 EXTRACT PREPARATION

Ethanol extract **EeC**; leaves were air-dried and ground then extracted with absolute ethanol, using cold maceration. The mixture was filtered and extract concentrated by evaporation, air dried and stored at 20°C till needed. For the aqueous extract **AeC;** leaves were air-dried and pulverised then macerated in distilled boiling water (100°C) and the marc obtained was left to stand for 24 hours and then filtered. The filtrate obtained was concentrated by evaporation at 50°C and the extract stored at 20°C.

### 2.2 PHYTOCHEMICAL ANALYSIS

Phytochemical studies were conducted on **AeC** and **EeC** for the presence or absence of Alkaloids, Flavonoids, Anthraquinones (free or combined), Tannins, Steroids, Saponins, Terpenoids, and Cardiac glycosides.

### 2.3 EXPERIMENTAL PROCEDURE

Male Wistar rats (8-10 weeks, weighing 130-170g) were used. All animals were housed under constant temperature and exposed to a 12hr light/12hr dark cycle and were fed a standard chow diet. Animal protocols were approved by the ethics committee for the care and use of laboratory animals at University of Lagos, Nigeria. After 1 week of acclimatization, the animals were randomly divided 5 per group for Aqueous and Ethanolic extracts into the following experimental groups;

A. Control Group received saline water
B. L-NAME only Group: received L-NAME (20mg/kg)

#### Aqueous extracts grouping; AeC

C. L-NAME + 100MG/KG

D. L-NAME + 200MG/KG

E. L-NAME + 300MG/KG

F.L-NAME + 400MG/KG

#### Ethanolic Extracts grouping; EeC

G. L-NAME + 100MG/KG

H. L-NAME + 200MG/KG

I. L-NAME + 300MG/KG

J. L-NAME + 400MG/KG

Blood pressure measurement for the duration of experiment was carried out with a power lab (TICE^®^).

Animals were anaesthetized humanely at the end of the experiments according to established protocols. The extracts and the standard agents were administered twenty minutes (20 mins) after L-NAME administration. Heparin solution was injected in the arterial catheter at intervals to avoid possible blood coagulation. Readings obtained were analyzed using the power lab and recorded. The safety effects of **AeC** and **EeC** were studied *In-vivo* on the systolic blood pressure, diastolic blood pressure and mean arterial blood pressure of rats treated with L-NAME.

### 2.4 ACUTE TOXICITY TEST

A single dose test was carried out for acute toxicity of the extracts. A dozen mice were fasted for 16 hours and then administered 5000mg/kg **AeC** and **EeC** separately. They were observed for 24 to 72 hours for signs of behavioural change and death.

### 2.5 STATISTICAL ANALYSIS

Statistical analysis was performed and carried out by student’s t-test using Graph Pad Prism 5.0. All results were expressed as means ± standard error of mean (SEM). Differences were considered significant at p <0.05.

## 3.0 RESULTS and DISCUSSION

Phytochemical analysis showed the presence of flavonoids, alkaloids, steroids, terpernoids and cardiac glycosides in the leaves of the test plant. Fig. 1.

The intravenous administration of **AeC** and **EeC** to L-NAME induced hypertensive rats caused a significant decrease in systolic blood pressure, diastolic blood pressure and Mean Arterial Blood Pressure. (Figures 2–5). Table 1 shows that the control groups were adequate as controls as the saline group did not show any significant increments or decrease in the parameters of importance while the L-NAME control group should an increase in the parameters. It is well known that L-NAME, administered either acutely by intravenous route or chronically by the oral means induce sustained hypertension (Nyadjeu *et al.*, 2013, Chaswal *et al.*, 2011).

**Figure 2 :**
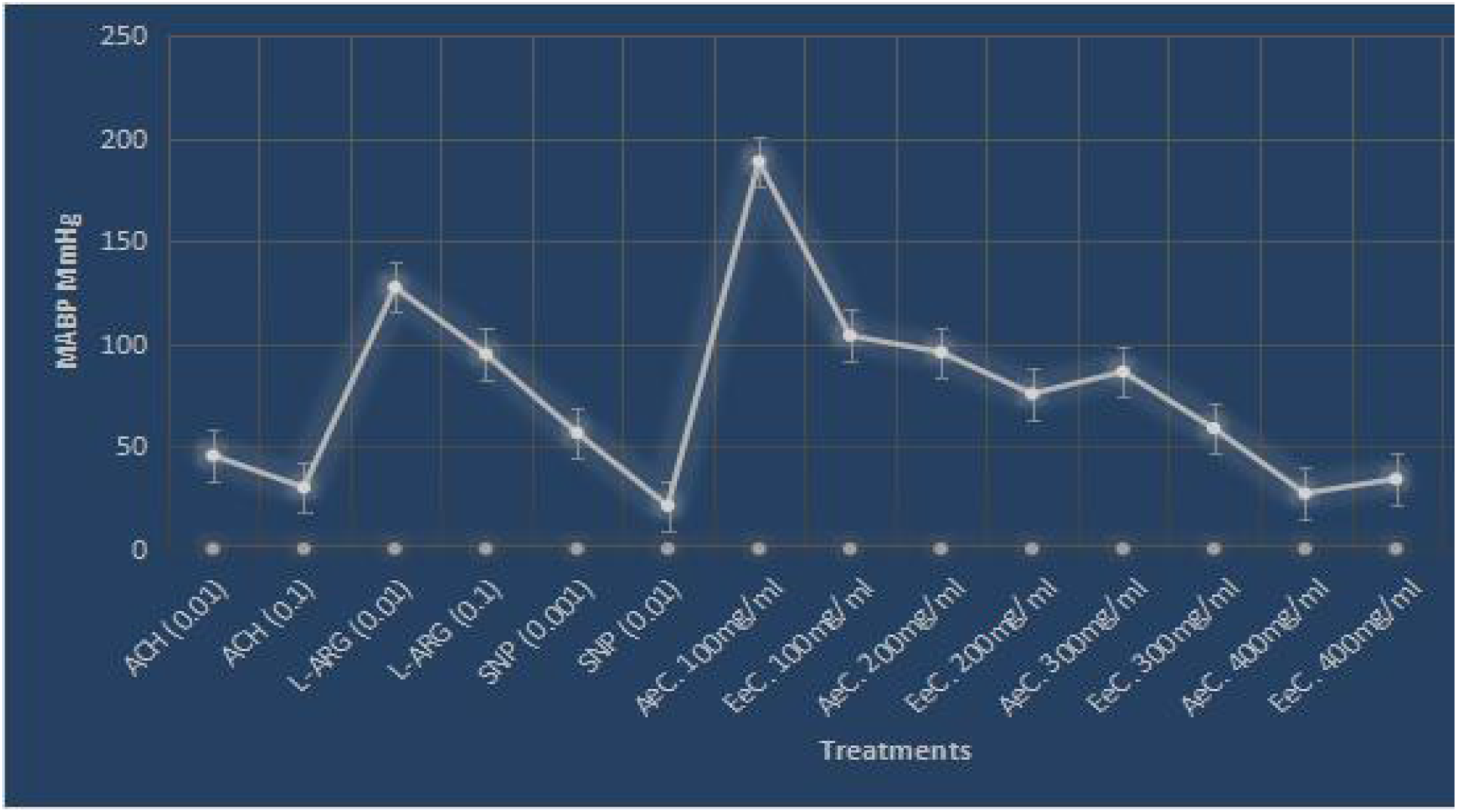
Effects of Standard drug agents and **AeC** and **EeC** on MABP. **Key:** ACH-Acetylcholine; L-ARG-L-Arginine; SNP- Sodium Nitroprusside (all in Moles). **AeC**- Aqueous extract; **EeC**- Ethanol extract.

**Table 1:**
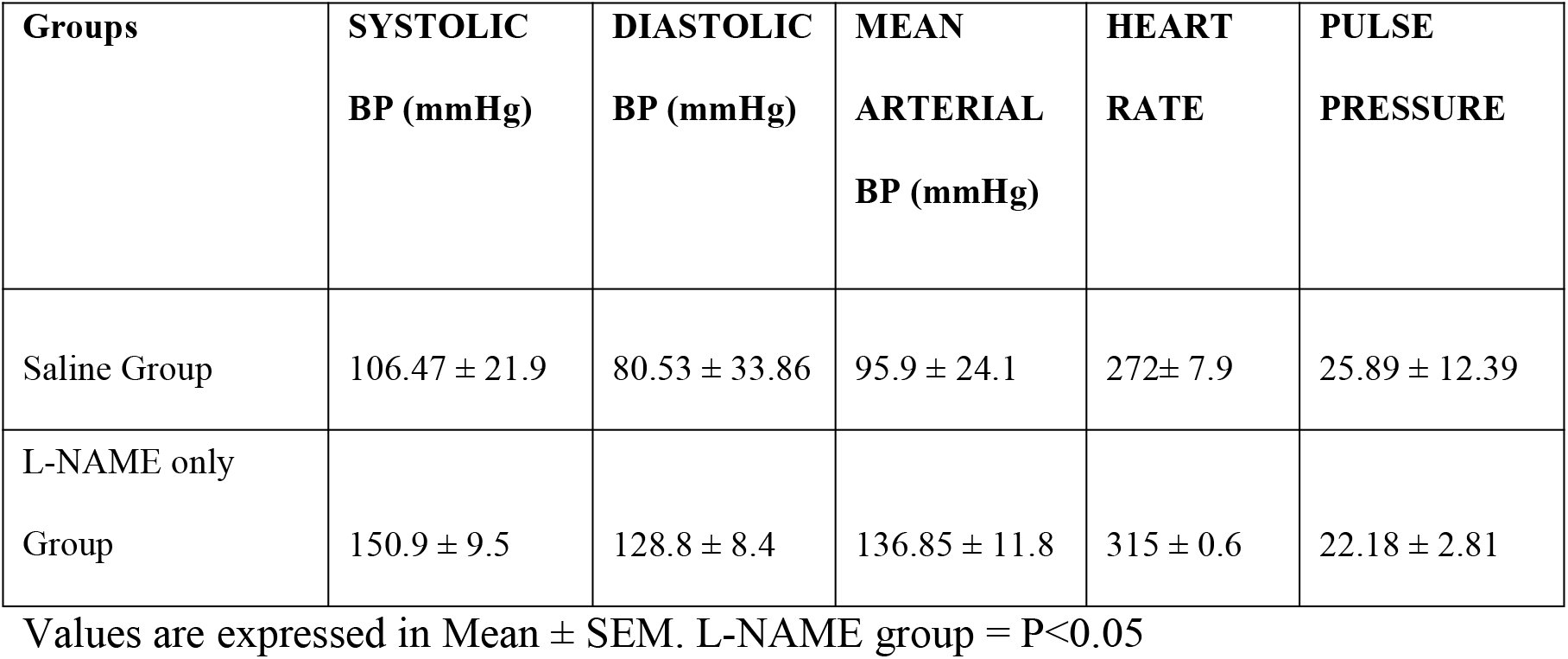
Effect of Saline and L-NAME on the Control animals.

| Groups | SYSTOLIC<br>BP (mmHg) | DIASTOLIC<br>BP (mmHg) | MEAN<br>ARTERIAL<br>BP (mmHg) | HEART<br>RATE | PULSE<br>PRESSURE |
| --- | --- | --- | --- | --- | --- |
| Saline Group | 106.47 ± 21.9 | 80.53 ± 33.86 | 95.9 ± 24.1 | 272± 7.9 | 25.89 ± 12.39 |
| L-NAME only<br>Group | 150.9 ± 9.5 | 128.8 ± 8.4 | 136.85 ± 11.8 | 315 ± 0.6 | 22.18 ± 2.81 |

Values are expressed in Mean ± SEM. L-NAME group = P<0.05

The genus *Clerodendrum* is well documented to have beneficial effects on the cardiovascular system of mammals due to its significant flavonoid content (Chan *et al.*, 2000; Arif-Ullah and Anwarul, 2008). Indeed, flavonoids from a great variety of plants have hypotensive and vaso-relaxant activities (Aduragbenro et al., 2009; Rosalia et al., 2010). Wang et al., (2018) conducted an extensive study of the diverse phytochemicals in aqueous and methanolic fractions of different plant parts and published them with relevant structures.

**AeC** at 400mg/ml appears to be the most effective dosage statistically (Figs.2-5) comparing very favourably with the standards ACH and SNP (Figs 2 and 3). L-NAME, when administered, induces a sustained mean arterial hypertension which persisted for over 30 minutes as observed here with a MABP of 136.85 ± 11.8 mmHg (Table 1). MABP was further elevated on administration of the lowest dose (100 mg/kg) of the **AeC** from 136.85 ± 11.8mmHg to 188.8 ± 0.8 mmHg. There was however a decline as the concentration increases from 95.8 ± 1.2mmHg to 86.3 ± 0.9mmHg respectively (Figure 2). The heart rate was also affected in Table 2, Fig. 4 and 5.

**Figure 3:**
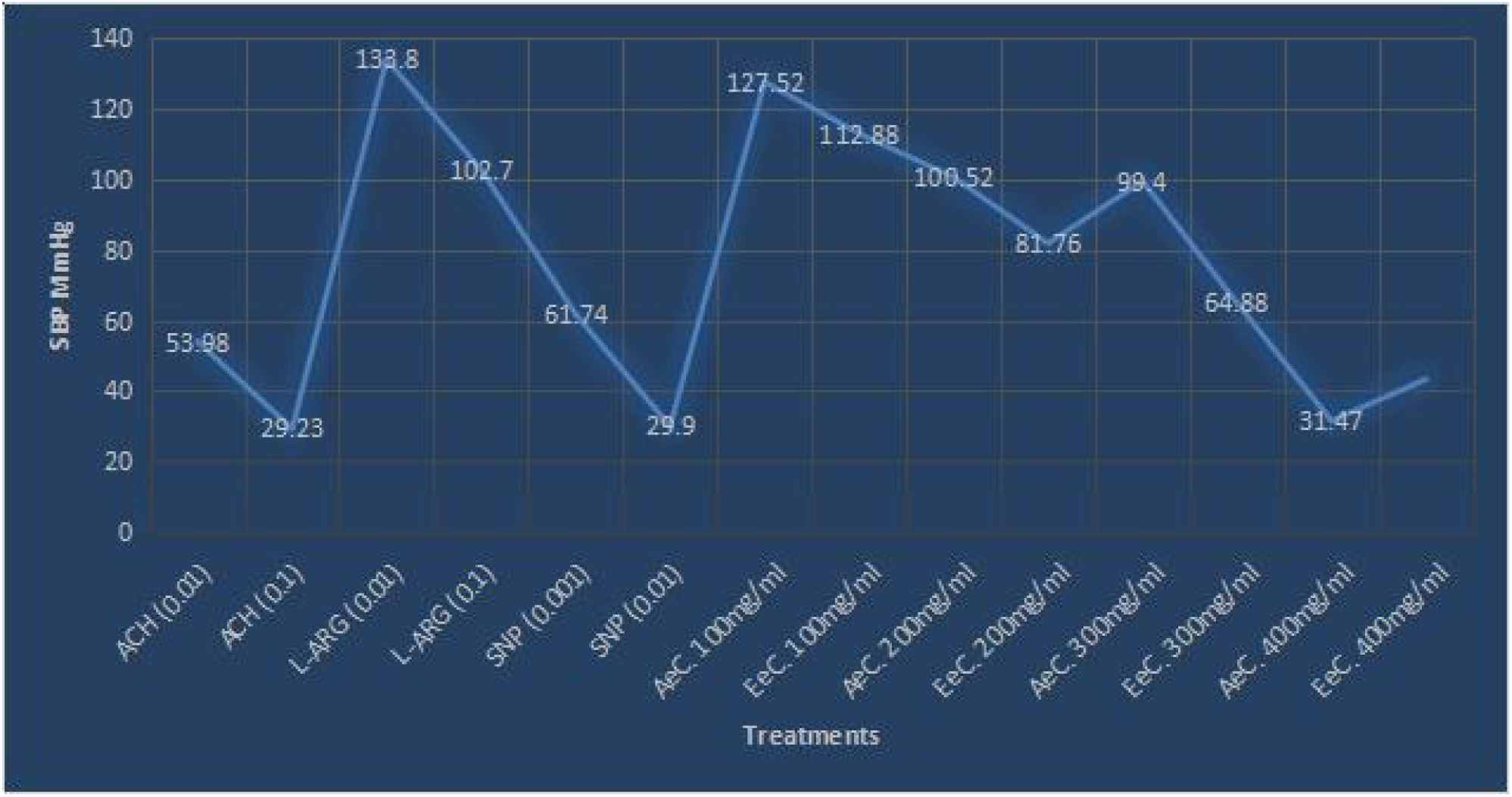
**E**ffects of Standard drug agents and **AeC** and **EeC** on SBP. **Key:** Standard Agents (P<0.01) - ACH-Acetylcholine; L-ARG- L-Arginine; SNP- Sodium Nitroprusside (all in Moles) **AeC**- Aqueous extract; **EeC**- Ethanol extract.

**Figure 4.**
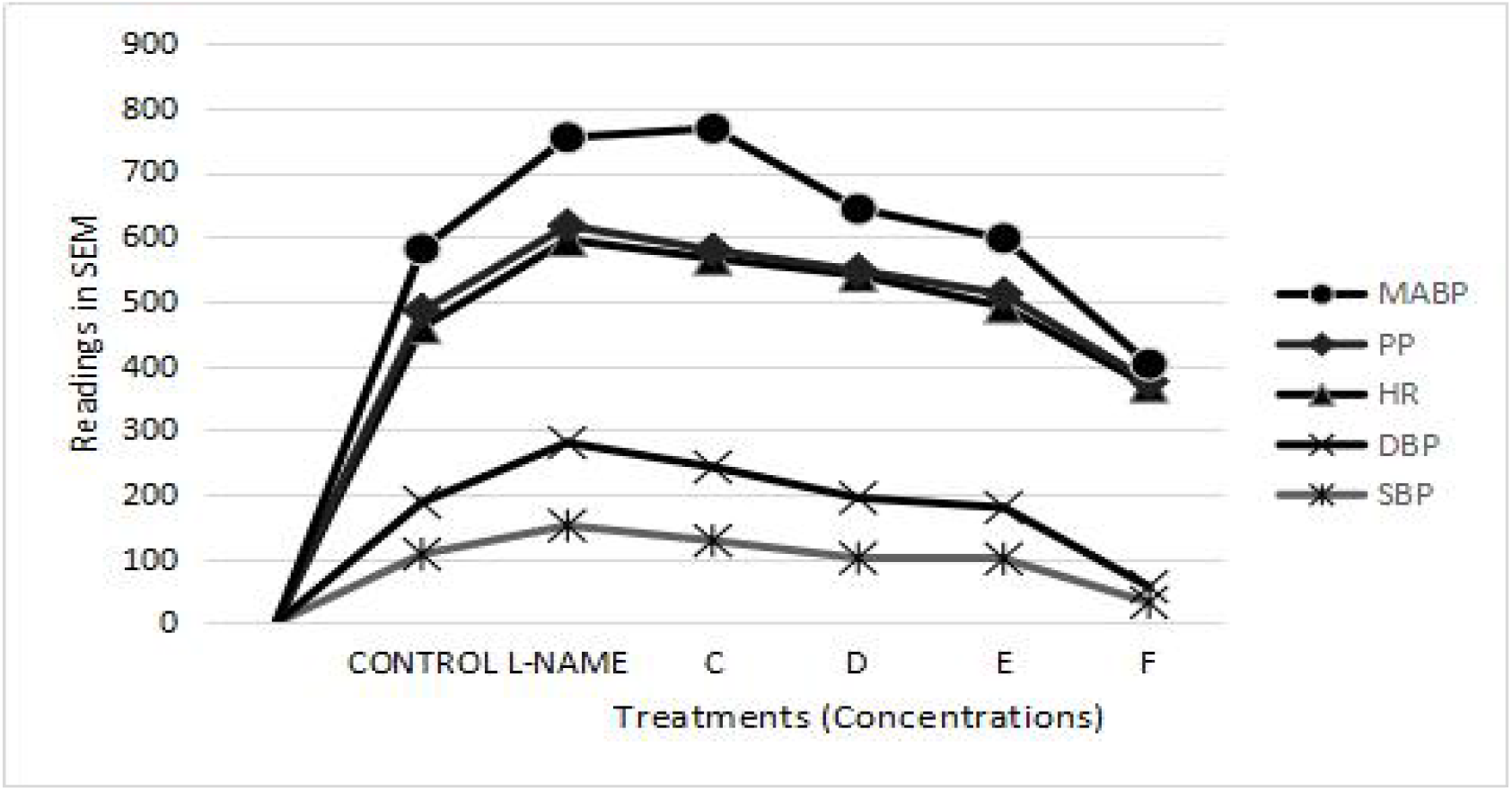
Effects of **AeC** on SBP, DBP, HR, PP and MABP

**Figure 5.**
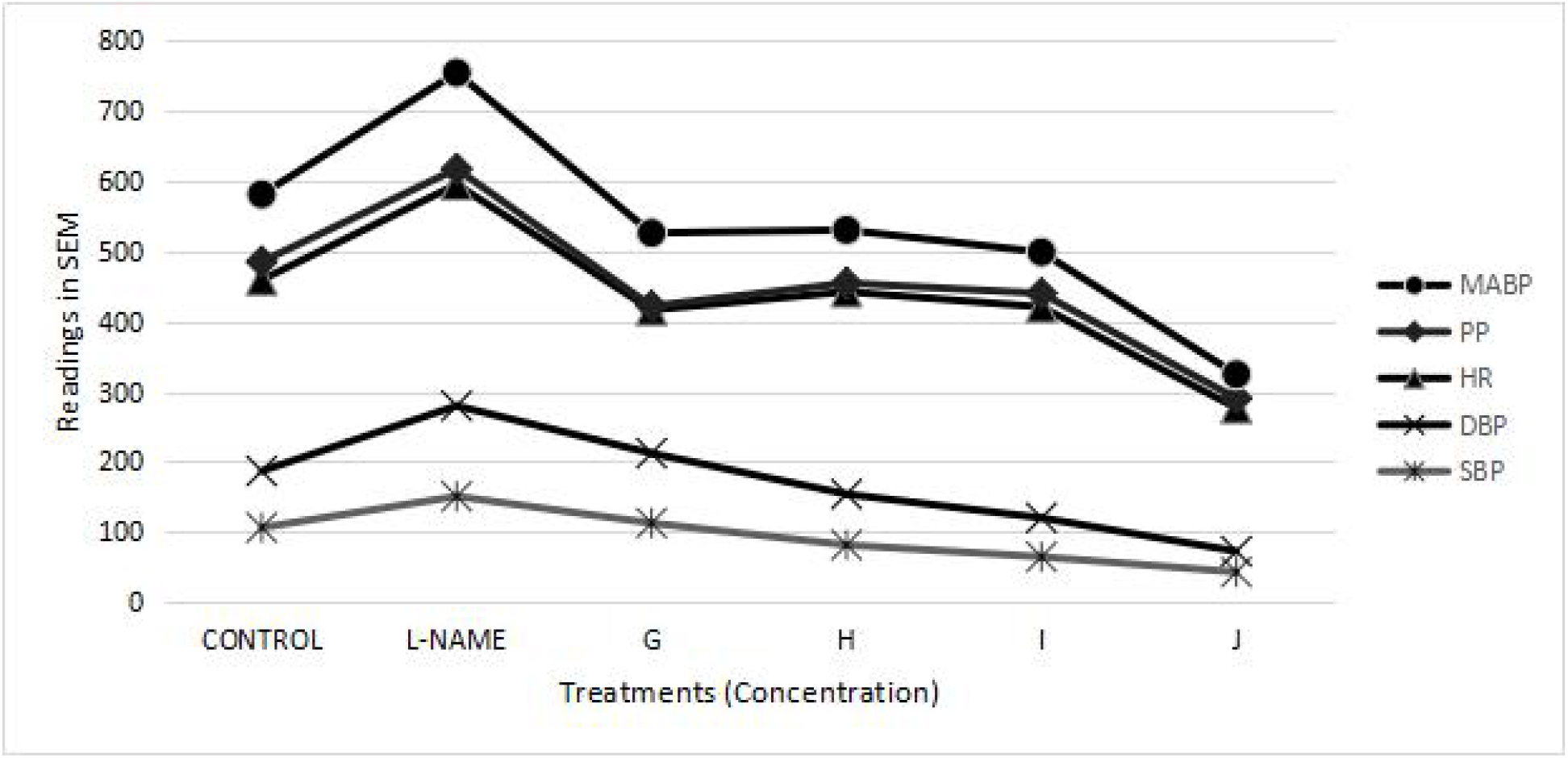
Effects of **EeC** on SBP, DBP, HR, PP and MABP.

**TABLE 2:**
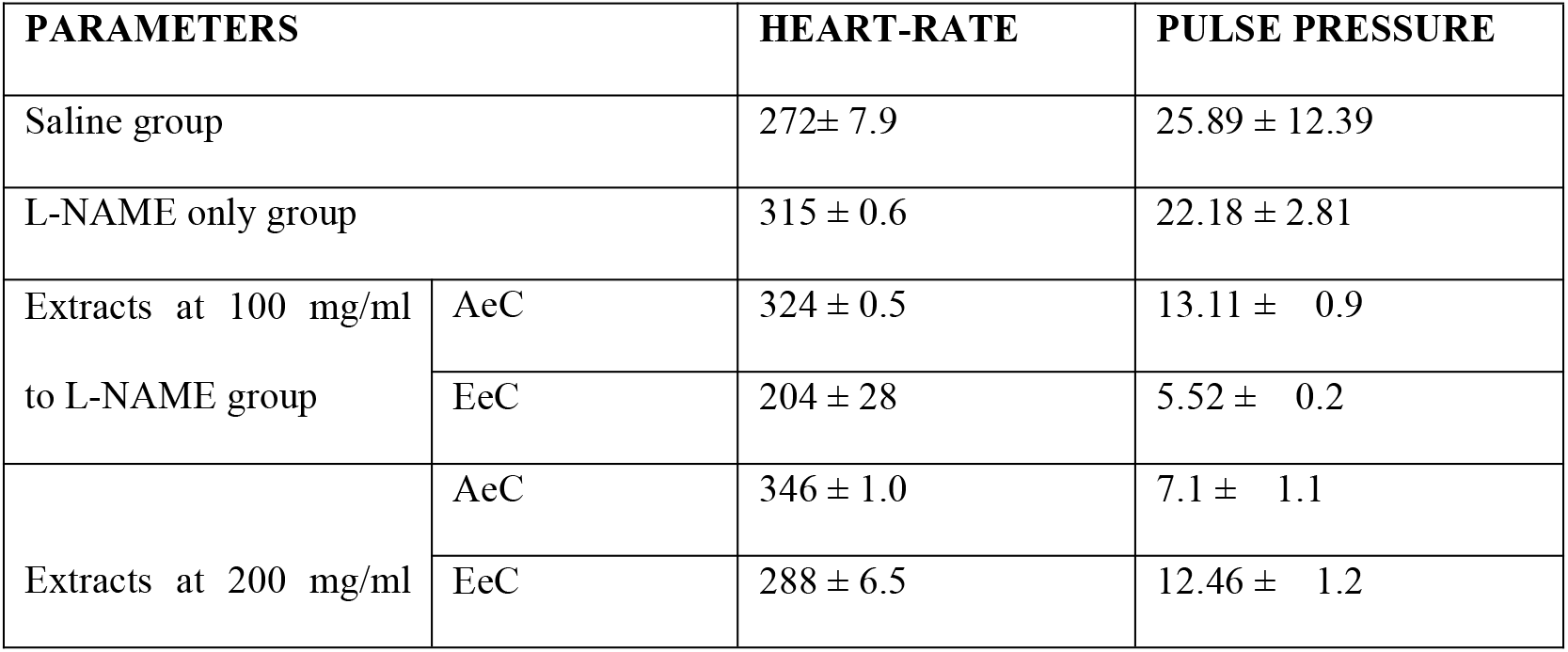
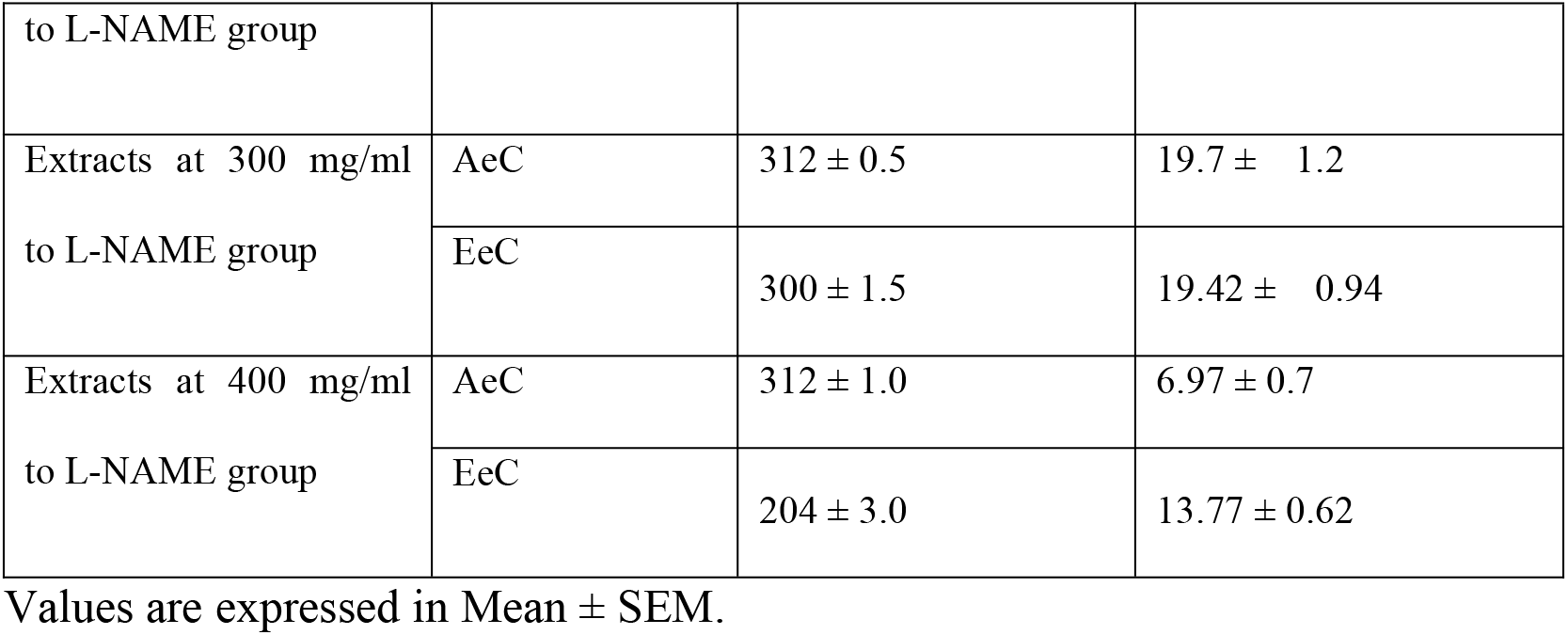
Effect of AeC and EeC on Heart rate and Pulse Pressure of L-NAME animals.

The MABP dropped suddenly after administration of 400mg/ml to 27± 1.5mmHg. The immediate fall in blood pressure induced by the doses (100, 200, 300 and 400 mg/ml) of the **AeC** was followed by a short-lasting effect indicating that the antihypertensive effects of the **AeC** are dose-related. The speed of the blood pressure drop is accounted for in local usage where the subjects are careful about the number of times in a day they partake of the decoction. 100mg/ml of the **EeC** reduced MABP to 103.9± 2.55 mmHg. 200mg/ml, 300mg/ml, 400mg/ml reduced the MABP to 75.5mmHg± 0.75, 58.7± 1.5, 34.1± 0.95mmHg respectively (Figure 2).

The effects were more lasting than that of the **AeC** and are observed to also be dose-related. The result, however, shows that the elevated MABP was reduced more by the **EeC** when compared to the **AeC** in the L-NAME induced hypertensive rats. **AeC**, at 100mg/ml, 200mg/ml and 300mg/ml reduced blood pressure and did not go below 90mmHg/ 60mmHg and can be considered as appropriate doses in reducing high blood pressure. For the **EeC**, only 100mg/kg was identified to reduce high blood pressure below 90mmHg/ 60mmHg and can be considered appropriate in reducing blood pressure also (Figures 3 and 4).

As in previous studies (Chaswal *et al.*, 2011), SNP administration leads to a rapid reduction in ABP (Figure 2) and when compared to both extracts, it can be said that SNP and the plants extracts successfully reduced the blood pressure (Figure 2). The effects of Acetylcholine, when compared to that of the extracts, was better but also shows a significant decrease in blood pressure by the extracts (Figures 2–4). Both extracts showed a significant decrease in blood pressure when compared to the standard agents (Table 2)

### 3.1 CONCLUSION

This study demonstrated the acute antihypertensive properties of the **AeC** and **EeC** in Nitric Oxide deficient hypertensive rats. The observed effects could partially be due to the reduction of the peripheral resistance by the thwarting the acute inhibitory effect of L-NAME on endothelial nitric Oxide Synthase. The recorded results authenticates its use in traditional medicine for the treatment of hypertension. *Clerodendrum chinensis* leaves extract, have shown to exhibit more potent antihypertensive properties when compared to known standards. Further studies are however required to check specific mechanisms of action and best and effective dosage forms.

## APPENDIX

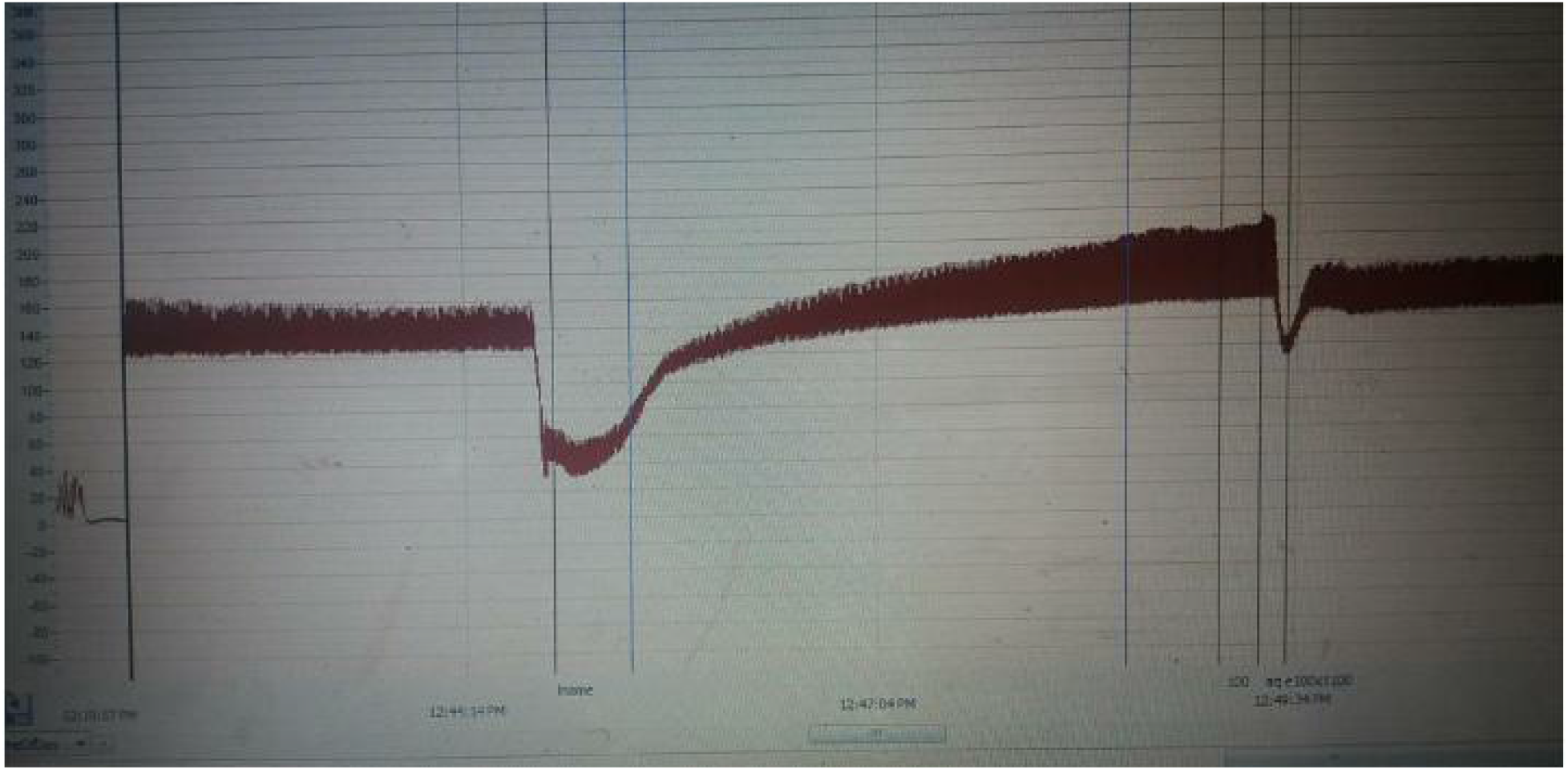
Peak readings for L-NAME

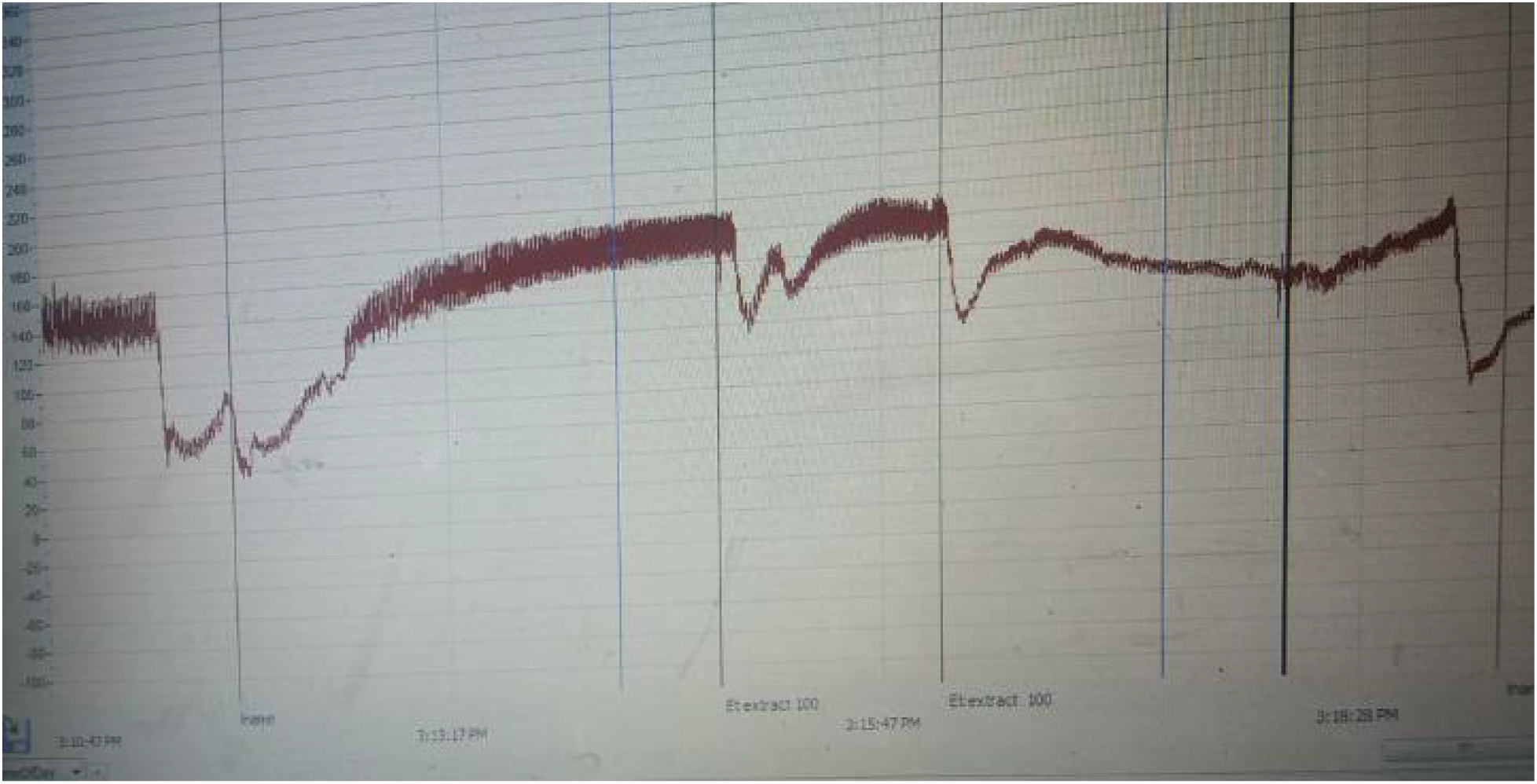
Peak readings for 100mg/kg **EeC**

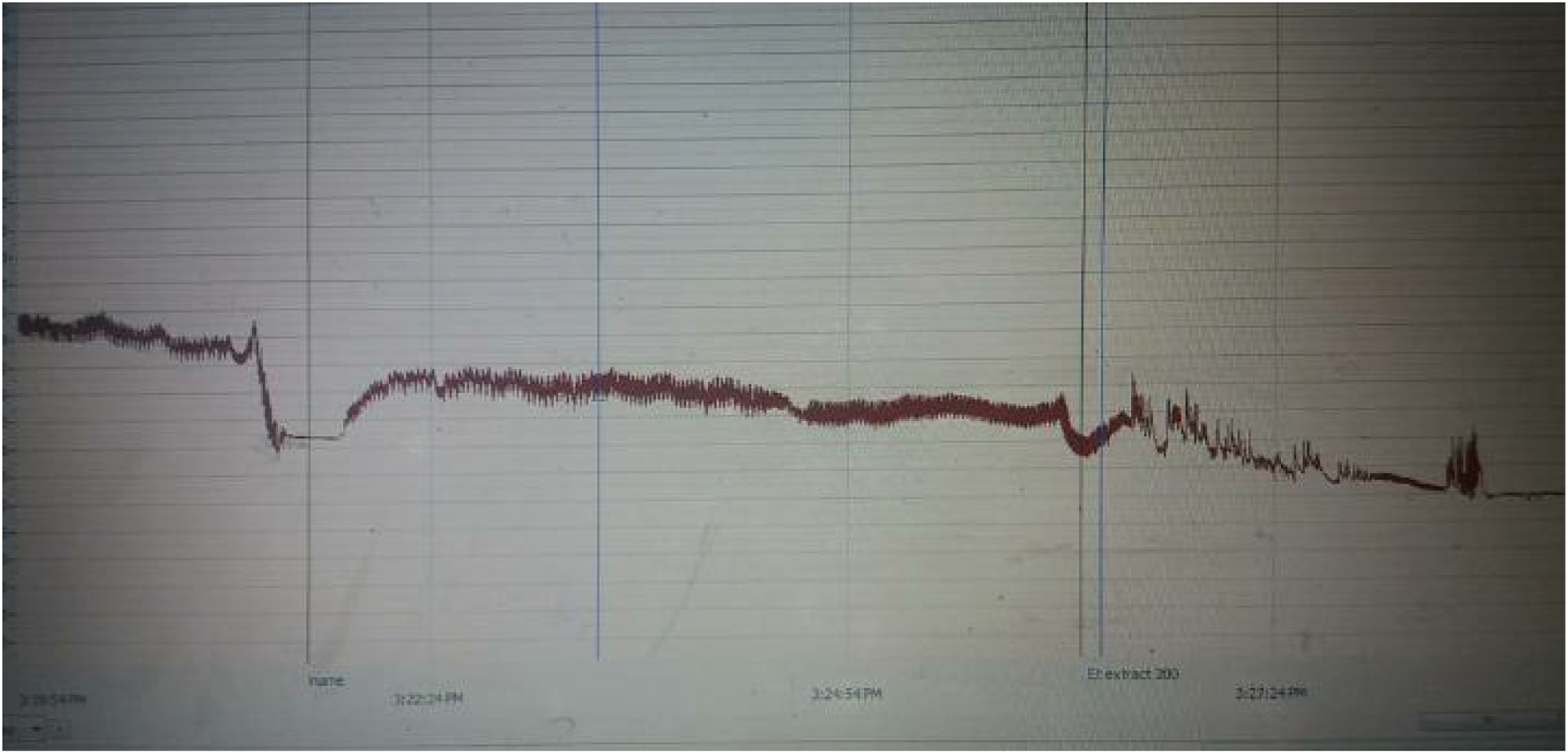
Peak readings for 200mg/kg **EeC**

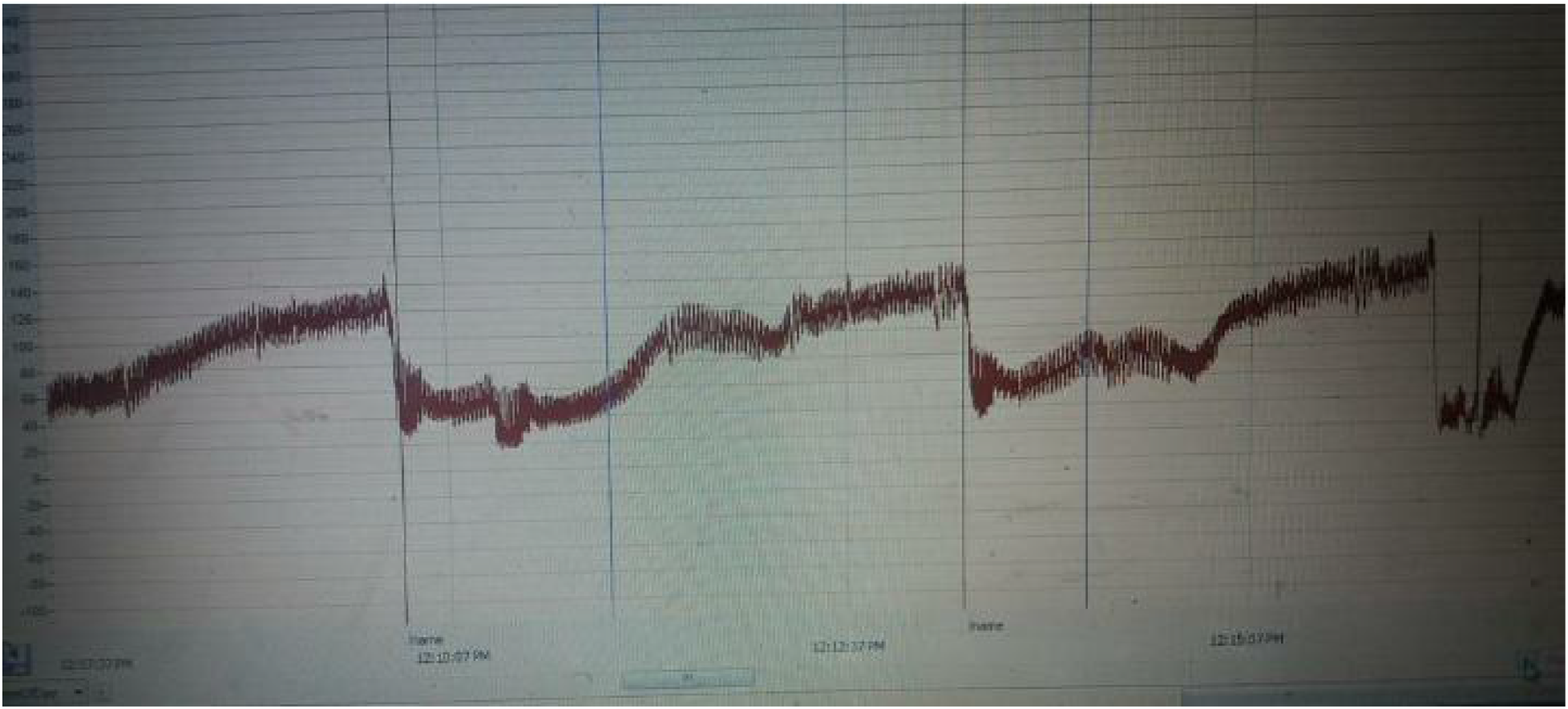
Peak readings for 300mg/kg **EeC**

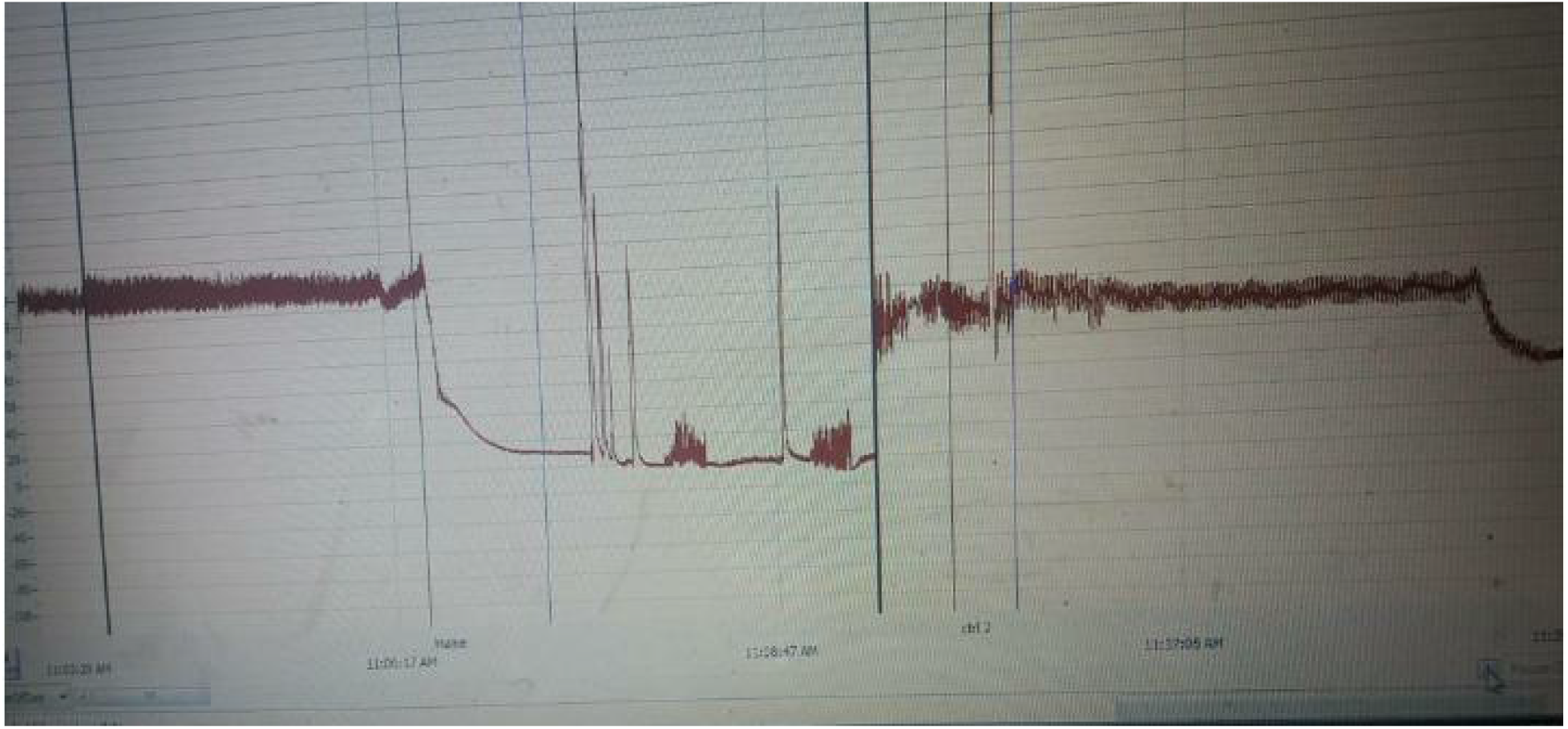
Peak readings for 400mg/kg **EeC**

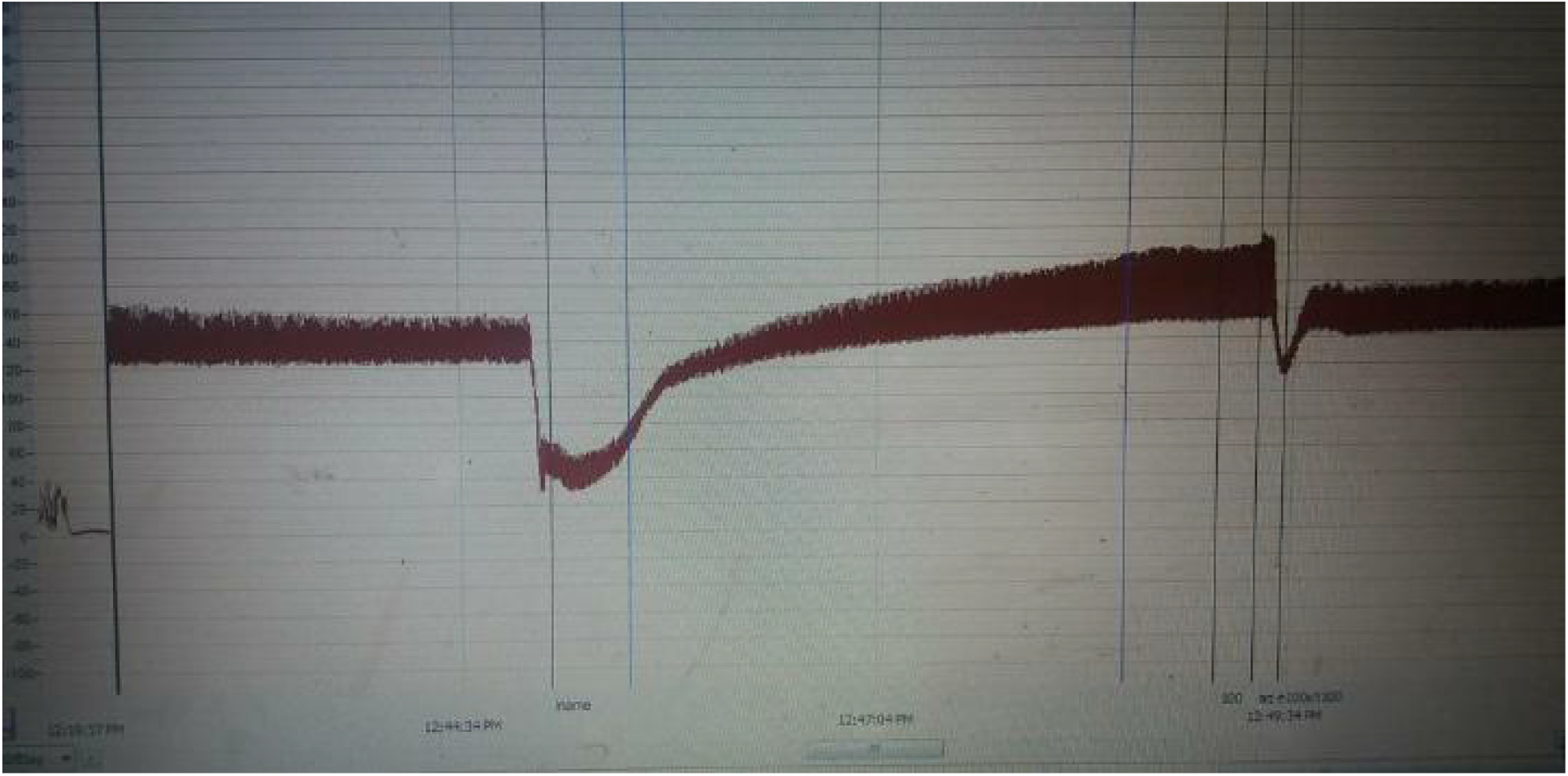
Peak readings for100mg/ kg **AeC**

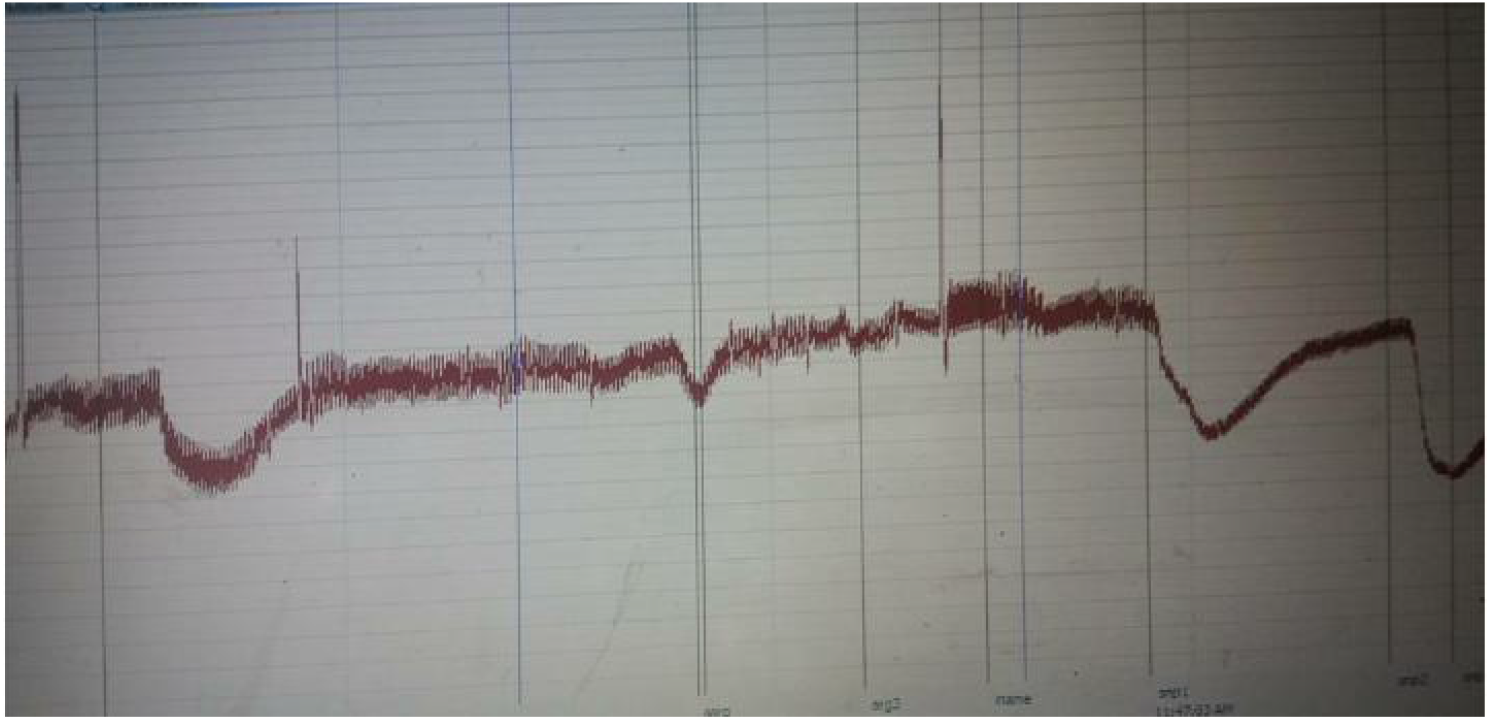
Peak readings for 200mg/ kg **AeC**

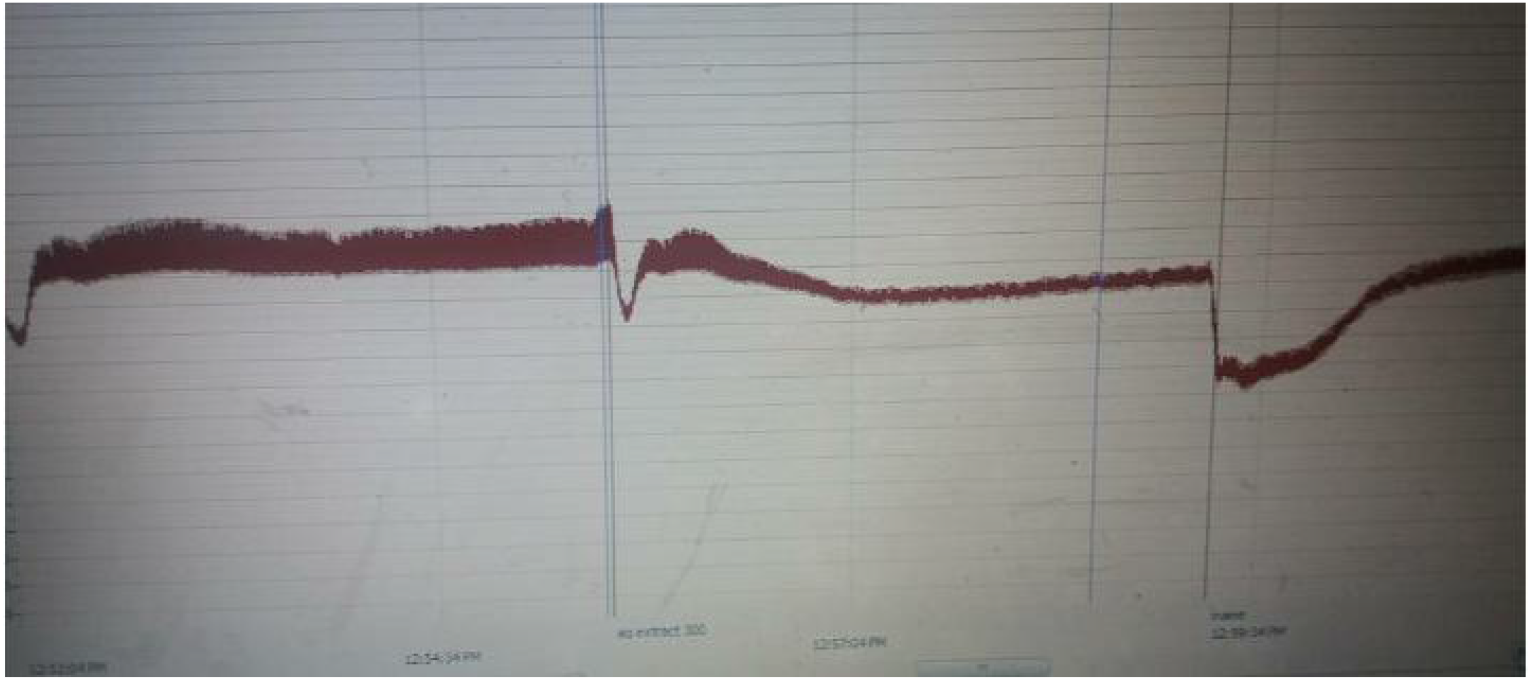
Peak readings for 300mg/ kg **AeC**

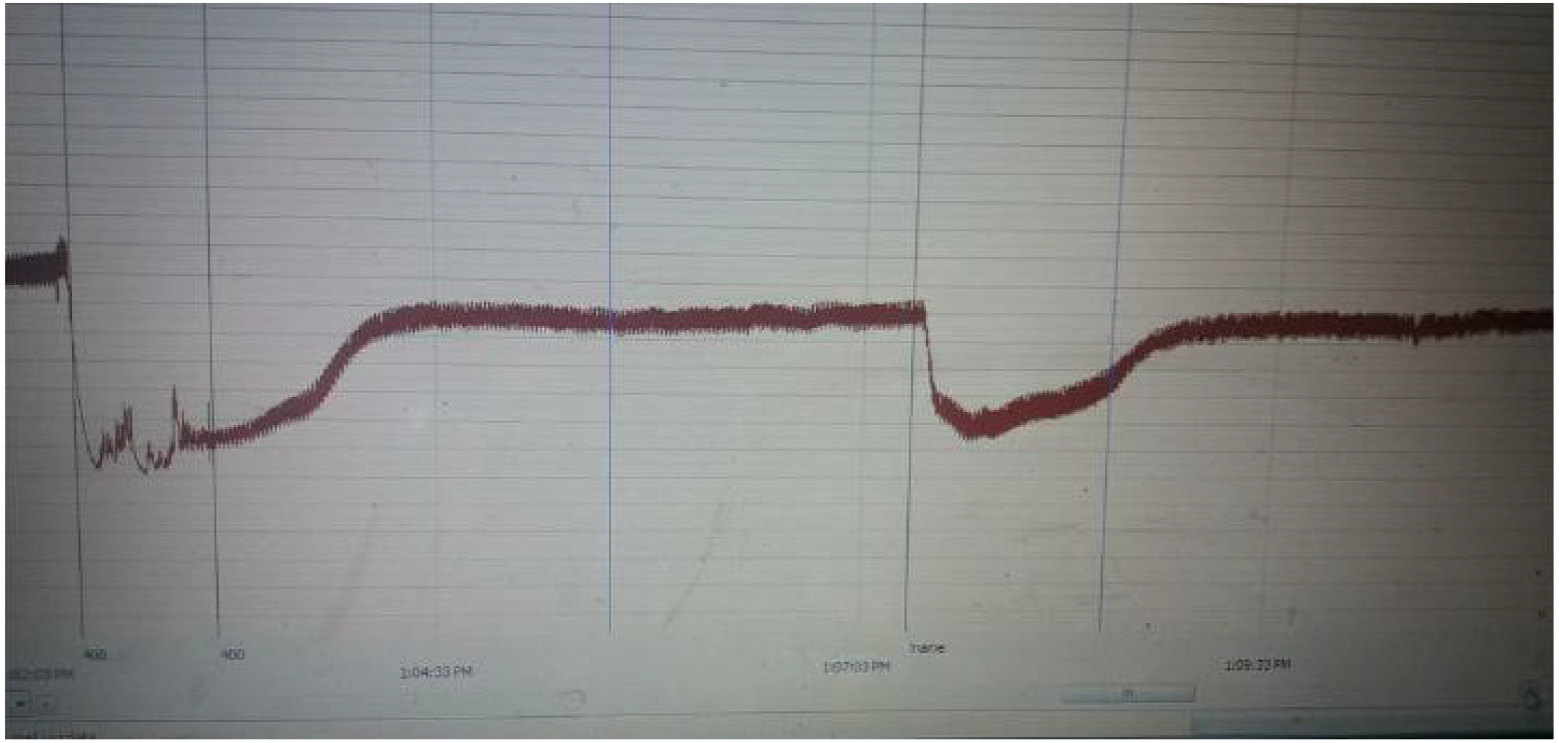
Peak readings for 400mg/ kg **AeC**

## Ethical Approval and Consent to participate

Not applicable

## Consent for publication

All the authors have seen the manuscript and agree to the publication

## Availability of supporting data

Not applicable

## Authors’ contributions

Odimegwu JI Conceptualized the research, designed the methodology, wrote up the original draft and corrected the final manuscript

Okanlawon, TF worked at Data curation and corrected Manuscript

Obumneme Noel NE carried out laboratory work and contributed to editing and review Ishola, IO supervised the laboratory work and data curation.

## Acknowledgements

We are grateful to Prof. C. Epie who collected the plant and told us about its actions and also to Mme R. Ossai for cultivating and multiplying the plant for more sample materials for this work.

